# Neuroblast sensory quiescence depends of vascular cytoneme contacts and sensory neuronal differentiation requires initiation of blood flow

**DOI:** 10.1101/667519

**Authors:** Laura Taberner, Aitor Bañón, Berta Alsina

**Affiliations:** Developmental Biology Unit, Department of Experimental and Health Sciences, Universitat Pompeu Fabra-Parc de Recerca Biomèdica de Barcelona, Dr. Aiguader 88, 08003 Barcelona, Spain

**Keywords:** sensory neurons, vascular, niche, cytoneme, Dll4/Notch, blood flow, zebrafish

## Abstract

In many organs, stem cell function depends on the communication with their niche partners. Cranial sensory neurons develop in close proximity to blood vessels, however whether vasculature is an integral component of their niches is yet unknown. Here, two separate, novel roles for vasculature in cranial sensory neurogenesis in zebrafish are uncovered. The first involves precise spatiotemporal endothelial-neuroblast cytoneme contacts and Dll4-Notch signalling to restrain neuroblast proliferation. Secondly, we find that blood flow onset triggers a transcriptional response to modify neuroblast metabolic status and is required for sensory neuron differentiation. In contrast, no role of sensory neurogenesis in vascular development is found, suggesting a unidirectional signalling from vasculature to sensory neuroblasts. Altogether, we demonstrate that the cranial vasculature constitutes a hitherto unrecognized niche component of the sensory ganglia that regulates the pace of their growth and differentiation dynamics.

**Highlights:**

♦ Vasculature is part of the cranial sensory ganglia niche and regulates neurogenesis.
♦ Cytoneme contacts between endothelial cells and sensory neuroblasts are required for neuroblast quiescence.
♦ Endothelial Dll4 and neuroblast Notch1 signal to regulate the growth of cranial sensory ganglia.
♦ Initiation of blood flow triggers a transcriptional metabolic switch and sensory neuronal differentiation.

## Introduction

Proper interaction and understanding of our external world depend on specialized sensory organs and neurons from the peripheral nervous system (PNS). The cranial PNS consists of several ganglia with sensory bipolar neurons that transmit external sensory information to the brain. Sensory neurogenesis is achieved by a coordinated sequence of specification, delamination, transit-amplification of progenitors and terminal differentiation events (Alsina et al., 2003; Begbie et al., 2002; Maier et al., 2014; Schlosser, 2010). Several intrinsic factors such as Igf1, FGFs or neurotrophins participate in the proliferation, innervation and function of cranial ganglia during embryogenesis and adulthood (Bilak et al., 2003; Camarero et al., 2003; Fantetti and Fekete, 2012; Murdoch and Roskams, 2012; Vemaraju et al., 2012). Mutations in some of these factors, therefore result in sensory pathologies, i.e. Igf1 mutations are associated with hearing loss in rodents and humans (Bonapace et al., 2003; Riquelme et al., 2010). The rate of sensory neurogenesis is determined by the balance between multipotent stem cells (SCs) and differentiated sensory neurons and the combination of intrinsic factors and extrinsic cues of their microenvironment.

Recent evidences indicate that multipotent SC function *in vivo* is tightly linked to communication with the components of the microenvironment in which they reside, the so called niche (reviewed in Spradling et al., 2001). Blood vessels have emerged as key participants of SC function in several niches, such as the adult neural, hematopoietic and spermatogonia SC niches. Depending on the context, blood vessels regulate SC quiescence, proliferation and/or differentiation (Palmer et al., 2000; Rafii et al., 2016; Shi and Gronthos, 2003; Yoshida et al., 2007).

The adult brain SC niche is one of the niches in which the blood vessel-neuronal interaction was early revealed and, thus, the signals and mechanisms are well understood. There, two types of neurovascular interactions take place, one mediated by direct cell-cell contacts through neural SCs (NSCs) endfeet enwrapping endothelial cells (ECs) (Kacem et al., 1998) and the other mediated by secreted signals from ECs (Shen et al., 2008; Tavazoie et al., 2008)(Mirzadeh et al., 2008; Palmer et al., 2000; Shen et al., 2008; Tavazoie et al., 2008). Cell-cell contact signalling by EphrinB2 and Jagged1 expressed in ECs and, Eph and Notch receptors in NSCs, regulates NSCs quiescence (Ottone et al., 2014). On the other hand, endothelia secrete VEGFs, Angiopoietins, BDNF, NT-3 and PEDF that control NSC self-renewal, progenitor cell proliferation or differentiation (Androutsellis-Theotokis et al., 2009; Calvo et al., 2011; Delgado et al., 2014; Jin et al., 2002a; Louissaint et al., 2002; Ramírez-Castillejo et al., 2006)(Androutsellis-Theotokis et al., 2009; Calvo et al., 2011; Delgado et al., 2014; Jin et al., 2002b; Louissaint et al., 2002; Ramírez-Castillejo et al., 2006). Blood vessels not only communicate through secreted factors, but also by the perfusion of oxygen and nutrients. Oxygen tension mediates the metabolic switch from glycolysis to Oxidative Phosphorylation (OxPhos), which has been shown recently to control NSC fate (Ito and Suda, 2014; Llorens-Bobadilla et al., 2015). In the embryo, vasculogenesis and neurogenesis are spatiotemporally coupled in the central nervous system (Hogan et al., 2004; Ulrich et al., 2011; Vasudevan et al., 2008), but only few studies have addressed the molecular mechanisms underlaying this neurovascular interaction (Lange et al., 2016; Tan et al., 2016; Tata et al., 2016).

Cranial ganglia in vertebrates lie in very close proximity to head blood vessels (Taberner et al., 2018), suggesting that vasculature might be an essential component of their niche. Notwithstanding, this question and the mechanisms involved in their communication remain to be addressed in the sensory PNS.

In this study, by taking advantage of an avascular model and double labelling of ECs and sensory neurons in zebrafish, we identify two distinct mechanisms by which vasculature regulates cranial sensory neurogenesis. *In vivo* imaging of the developing sensory ganglia reveals that ECs and neuroblasts establish direct contacts through filopodial protrusions. Inhibition of the filopodial contacts show that they exert signalling functions over sensory neuroblast. Thus, the work establishes first evidences for neurovascular interaction mediated by cytonemes. Genetic and pharmacological perturbations show that cytoneme contacts and Dll4 signalling inhibit neuroblasts cell cycle entry and regulate quiescence at early timepoints. Later, blood flow initiation activates a transcriptional program linked to oxygen sensing by Egln2 and a metabolic switch to OxPhos, together with the promotion of cranial ganglia differentiation. Our results uncover, for the first time, that the cranial vasculature is an essential component of the sensory ganglia niche and can temporally regulate neuroblast quiescence and differentiation by using distinct signalling mechanisms.

## Results

### Vasculature regulates the number of SAG neuroblasts by restraining cell cycle entry of neuroblasts

We focused on neurovascular interactions in the statoacoustic ganglion (SAG), which comprise the sensory neurons that innervate mechanosensory hair cells of the inner ear and the brainstem. During neurogenesis, *neurog1*^+^ neuronal precursors (blue) transit into neurod^+^ neuroblasts (green) that delaminate and coalesce into the SAG. Later they differentiate into Islet2^+^ neurons (orange) (Figure 1A). The developing zebrafish SAG lies between three main vessels, the Primordial Hindbrain Channel (PHBC) that runs between the hindbrain and the otic vesicle (OV), the Lateral Dorsal Aorta (LDA) beneath the OV and the Primary Head Sinus (PHS) surrounding laterally the OV (Taberner et al., 2018) (Figure 1B).

**Figure 1.**
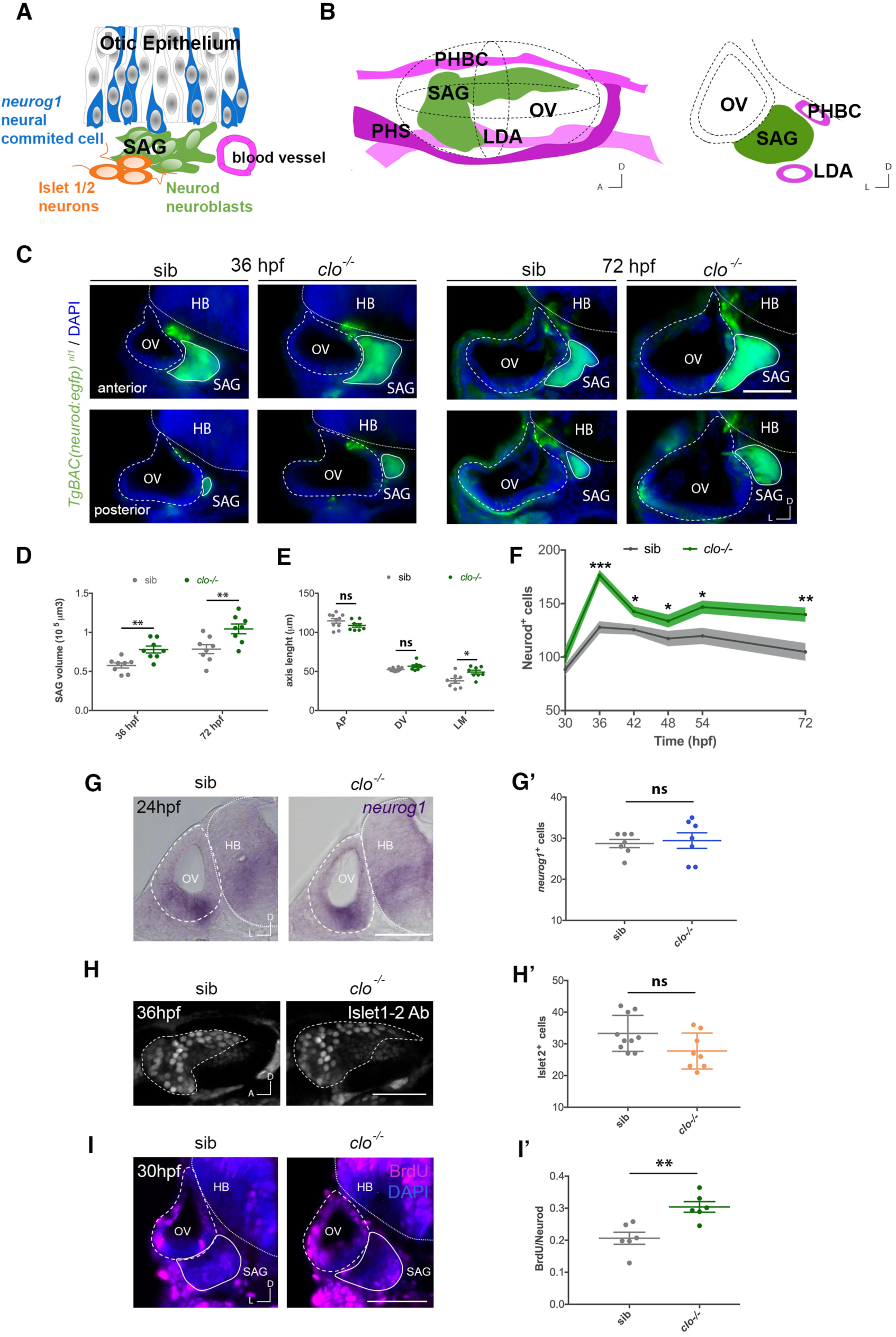
In the absence of vessels, the Neurod^+^ population in the SAG is expanded due to increased proliferation but not to effects on specification or differentiation. (A) Schematic drawing of the sequential steps of otic neurogenesis. Coloured graphs representing effects in specific cell types will follow this scheme: neurog1^+^ neuronal committed progenitors in blue, Neurod^+^ neuroblasts in green, Islet1/2^+^ neurons in orange and blood vessels in magenta. (B) Lateral (left) and transversal (right) 3D drawing of the OV (dotted line), the SAG and the adjacent vessels: PHBC, LDA and PHS. (C) Representative anterior and posterior transverse images of the SAG in sib and clo^-/-^ *TgBAC(neurod:egfp)* ^*nl1*^ embryos of cryostat sections, at 36 and 72 hpf. Neurod^+^ cells: green, neuroblasts; DAPI: blue, nuclei. (D) Quantification of SAG volumes in sib and clo^-/-^ embryos at 36 and 72 hpf (n=8). (E) Graph showing measurements of the anteroposterior (AP), dorsoventral (DV) or mediolateral (ML) axis length of the SAG in sib and clo^-/-^ embryos (n=8-10), at 72 hpf. (F) Quantification of Neurod^+^ nuclei in the SAG visualized through the *TgBAC(neurod:egfp)* ^*nl1*^ of sib and clo^-/-^ embryos at different timepoints from 30 to 72 hpf (n = 6-10). (G) Transversal images of *neurog1 in situ* hybridization in the OV epithelium in sib and clo^-/-^ embryos, at 24 hpf; and quantified in G’ (n = 7). (H) Confocal lateral images of immunostained Islet 1/2^+^ nuclei in the SAG (dotted line, visualized by *TgBAC(neurod:egfp)* ^*nl1*^) in sib and clo^-/-^ embryos, at 36 hpf; and quantified in H’ (sib n = 10, clo^-/-^ n = 8). (I) Transversal images of BrdU incorporation (magenta) in the SAG (continuous line, visualized by *TgBAC(neurod:egfp)* ^*nl1*^) in sib and clo^-/-^ embryos, at 30 hpf; and quantified in I’ (n = 6). DAPI: blue, nuclei. Scale bar, 50 µm. Error bars, mean ± SEM for D,E,G’,H’,I’; and mean ± SD for F. Unpaired two-tailed Student’s *t*-test. * p < 0.05, ** p < 0.01, *** p < 0.001. ns, non-significant. OV, otic vesicle; SAG, statoacoustic ganglion; PHBC, primordial hindbrain channel; LDA, lateral dorsal aorta; PHS, primary head sinus; HB, hindbrain; sib, siblings.

To functionally evaluate if cranial vasculature regulates the development of cranial sensory neurons in SAG, we analysed SAG growth dynamics in *cloche* (*clo*) homozygous mutants in the double *TgBAC(neurod:EGFP)* ^*nl1*^;*(Kdrl:ras-mCherry)* ^*s896*^ background, in which eGFP marks sensory neurons and mCherry marks ECs. *Cloche* encodes *npas4l*, a PAS-domain-containing bHLH transcription factor required for the formation of endothelial and hematopoietic cells (Reischauer et al., 2016; Stainier et al., 1995) and *clo* mutants have extensively been used as an avascular model (Liao et al., 1997; Sumanas et al., 2005). In *clo* mutant embryos, the SAG appeared larger compared to siblings (sib) at 36 and 72 hpf (Figure 1C). Quantification of SAG volumes revealed a statistically significant increase in *clo* mutant embryos (Figure 1D). Moreover, the SAG displayed an abnormal shape and a significant expansion along its mediolateral axis but not along the anteroposterior or dorsoventral axis (Figures 1C and 1E). The increased volume in *clo* mutant embryos could reflect a loss of confinement of the SAG cells by taking the space left by the lack of blood vessels or, alternatively, a higher number of Neurod^+^ cells. To distinguish between both possibilities, we quantified in serial transverse sections the nuclei within the GFP-labelled SAG cells between 30 to 72 hpf in the *clo* mutant and sibling embryos (Figures 1F and S1). In siblings, there is a period of steep SAG growth between 30 to 36 hpf, (sib 30 hpf: 89 ± 5; sib 36 hpf: 128 ± 5). From 36 to 72 hpf, the mean number of Neurod^+^ cells is 105 ± 8, suggesting that after 36 hpf this population does not grow. Remarkably, we found that the number of Neurod^+^ cells was significantly higher in *clo* mutant embryos compared to siblings at 36 hpf (sib: 128 ± 5 vs *clo*^-/-^: 177 ± 5) and this increase persisted at all the following timepoints (Figure 1F). Several mechanisms could account for the increased number of Neurod^+^ cells in *clo* mutant embryos: i) increased specification of *neurog1*^*+*^ cells within the otic epithelium and thereby increased delamination of Neurod^+^ cells, ii) reduced differentiation of Neurod^+^ cells into Islet2^+^ neurons or iii) increased proliferation of the transit-amplifying Neurod^+^ population. We did not find a higher number of *neurog1*^*+*^*-*expressing cells in the otic epithelium in *clo* mutant embryos compared to siblings (Figures 1G and 1G’) nor significant differences in the number of differentiated Islet2^+^ cells in the SAG (Figures 1H and 1H’). Instead, in *clo* mutant embryos we detected a significantly higher number of proliferating Neurod^+^ cells after a BrdU pulse from 24 to 30 hpf (expressed as BrdU^+^/Neurod^*+*^ ratio) (Figures 1I and 1I’).

Hence, our results show that at early timepoints, cranial vasculature regulates the number of SAG neuroblasts by negatively controlling cell proliferation of Neurod^+^ cells and therefore constitutes an essential component of the neurosensory niche during development.

### Cranial blood vessels and sensory neuroblasts establish direct contacts through filopodia

In both the embryonic and adult brain, NSCs establish contacts with ECs through endfeet projections (Ottone et al., 2014; Tan et al., 2016; Tavazoie et al., 2008). Therefore, we wondered if neuroblasts from the SAG also establish direct contacts with the adjacent vessels. To investigate this in detail, we live-imaged sensory neuroblasts from the SAG and ECs from the PHBC in double *TgBAC(neurod:EGFP)* ^*nl1*^;*(Kdrl:ras-mCherry)* ^*s896*^ reporter lines, at high spatiotemporal resolution. At 30 hpf, we did not observe neuronal extensions enwrapping ECs of the PHBC, but instead observed dynamic, thin filopodia extending from ECs to Neurod^+^ cells, as well as similar filopodia from Neurod^+^ cells extending to ECs (Figures 2A, 2B, S2, S3 and Supplemental Movies S1-4). The dynamics of filopodia were varied; in some instances, we observed filopodia extending and retracting in seconds (Figures 2A and 2B, cyan and magenta arrowheads respectively), in others, filopodia remained stable for several minutes (Figures 2A and 2B, yellow arrowheads). Interestingly, some PHBC filopodia did not remain at the surface of the SAG but contacted deeper neuroblast somas (Supplemental Movie S4). Moreover, stable filopodia-filopodia contacts between neuroblasts and ECs were also seen in some cases (Figures 2C and S3). As observed in transverse optical confocal reconstructions of the vessels and the SAG, filopodia were highly directional: PHBC filopodia extended only ventrally in the direction of sensory Neurod^+^ cells (Figures 2D and 2E). Interestingly, filopodia were present from 24 to 36 hpf but not later, suggesting an early role of filopodia in vessel-neuroblast communication (Figure 2F). Altogether, highly directed and temporally regulated filopodia-mediated cell contacts are established between both EC and sensory neuroblasts.

**Figure 2.**
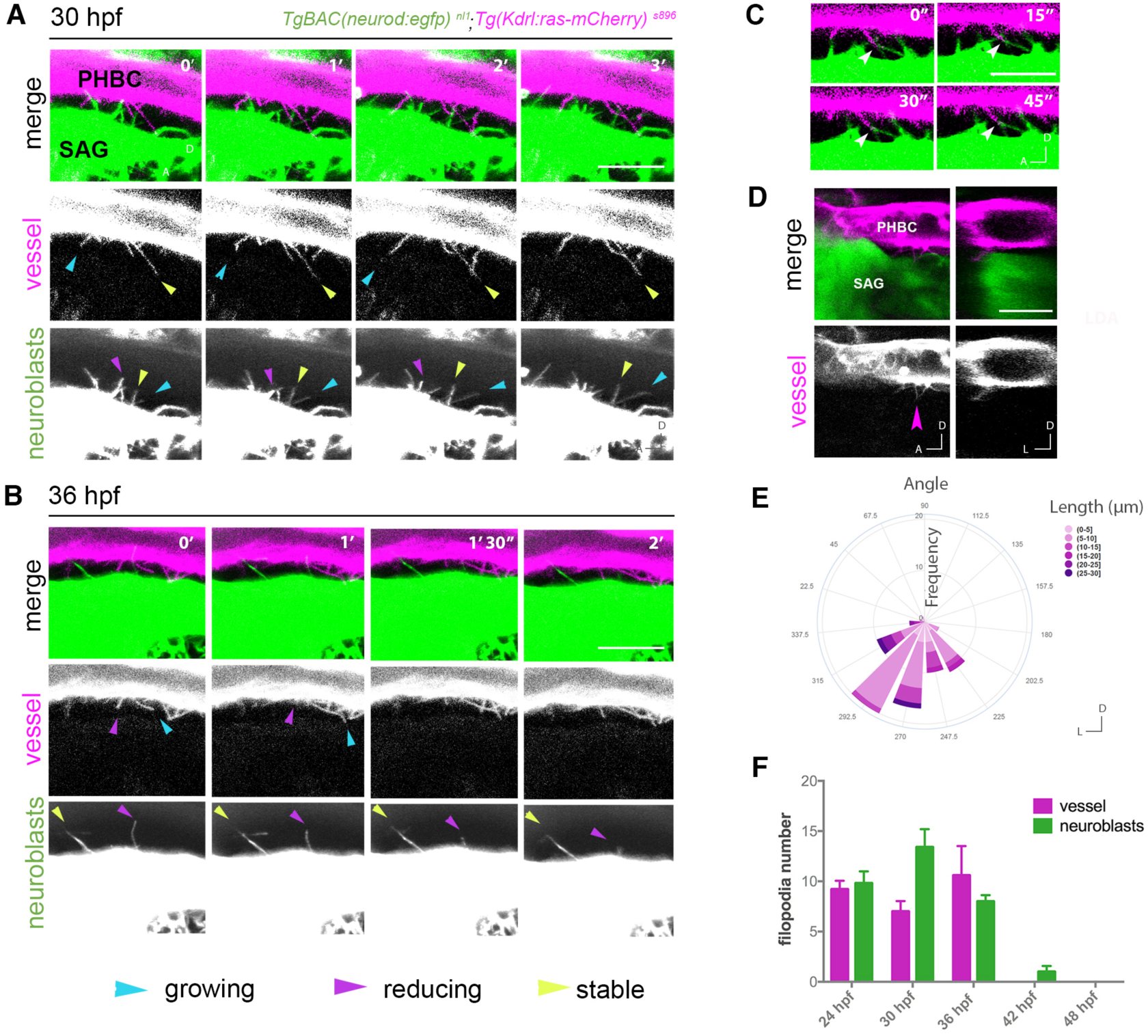
Endothelial cells and SAG neuroblasts contact through directional filopodia. (A,B) Timelapse of lateral images of *TgBAC(neurod:egfp)* ^*nl1*^; *(Kdrl:ras-mCherry)* ^*s896*^ embryos showing filopodia extending from the PHBC to the SAG and from the SAG to the PHBC at 30 and 36 hpf (videos are representative of 5 experiments). Cyan, magenta and yellow arrowheads point to examples of growing, reducing and stable filopodia, respectively. (C) 4 µm thick z-projection exemplifying a filopodia-filopodia contact remaining stable for more than 45 seconds, 30 hpf. (D) Single plane images in lateral view and transversal optic reconstruction showing a filopodial process from the PHBC to the SAG, at 30 hpf. Arrowhead indicates point for transverse optical reconstruction. (E) Rose plot of PHBC filopodia frequency of angle distribution and length (0°, lateral; 90°, dorsal; 180°, medial; 270°, ventral (n=76 filopodia, 15 embryos), at 30 hpf. (F) Histogram showing the number of filopodia in the PHBC (magenta) and the SAG (green) at different timepoints from 24 to 48 hpf (n = 5). Scale bars, 20 µm in A,B,D and 10 µm in C. SAG, statoacoustic ganglion; PHBC, primordial hindbrain channel.

### Sensory neurons are not required for cranial vascular patterning

Communication between neurons and vessels in the CNS can take place from vessels to neurons, vice-versa and bidirectionally (Barber et al., 2018; Makita et al., 2008; Mukouyama et al., 2002). Thus, we next investigated if sensory neuron development has a role in vascular patterning near sensory ganglia. In this case, we analysed homozygous *neurog1*^*hi1059*^ mutant embryos -which fail to specify sensory neuronal precursors (Ma et al., 1998)-in the double transgenic *TgBAC(neurod:EGFP)* ^*nl1*^;*(Kdrl:ras-mCherry)* ^*s896*^ reporter line. Strikingly, despite the total absence of SAG and other cranial ganglia in *neurog1* mutant embryos, development of the PHBC and LDA displayed no obvious patterning defects at 30 hpf (Figure 3A). The central arteries (CtAs) ingrowth into the hindbrain also did not display any delay nor obvious alteration at 48 hpf (Figure 3A). We quantified the width of PHBC and the number of ECs at 48 hpf, and again found no significant differences between *neurog1* mutants and siblings (Figures 3B and 3C). In conclusion, our data indicate that cranial vascular development does not require signals from sensory neuroblasts and suggest that primarily the signalling direction goes from vessels to neurons.

**Figure 3.**
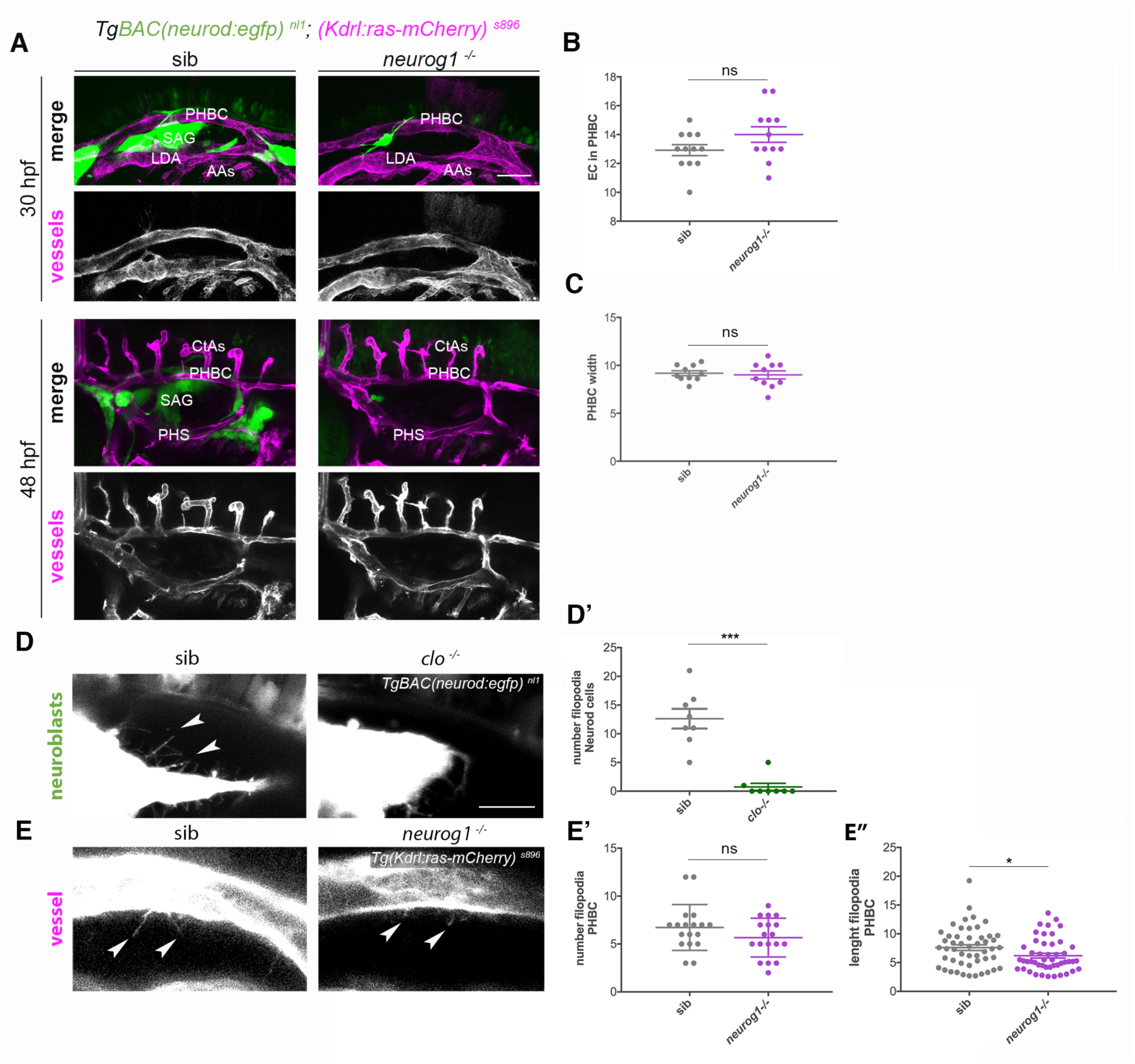
Lack of neurogenesis does not affect cranial vasculature patterning nor filopodia formation, while loss of vasculature prevents neuroblasts filopodia formation. (A) Representative lateral images of cranial vasculature in sib and *neurog1*^*-/-*^ homozygous mutant embryos in *TgBAC(neurod:egfp)* ^*nls1*^, *(Kdrl:ras-mCherry)* ^*s896*^ background, at 30 hpf and 48 hpf. (B) Quantification of EC numbers in sib and *neurog1*^*-/-*^ (n = 12), at 48 hpf. (C) Quantification of PHBC width in sib and *neurog1*^*-/-*^ (n = 10), at 48 hpf. (D) Confocal lateral images showing neuroblasts filopodial processes visualized through the *TgBAC(neurod:egfp)* ^*nl1*^ in sib and clo ^*-/-*^, at 30 hpf. (D’) Graph of SAG neuroblasts filopodia number in sib and clo ^*-/-*^ (n = 8), at 30 hpf. (E) Confocal lateral images showing PHBC filopodial processes visualized through the *Tg(Kdrl:ras-mCherry)* ^*s896*^ in sib and *neurog1*^*-/-*^ embryos, at 30 hpf. Arrowheads indicate filopodia in D and E. (E’) Graph of PHBC filopodia number in sib and *neurog1*^*-/-*^ (n = 18), at 30 hpf. (E’’) Graph of PHBC filopodia length in sib (n = 50) and *neurog1*^*-/-*^ (n = 48), at 30 hpf. Scale bars, 50 µm in A and 20 µm in D,E. Error bars, mean ± SEM. Unpaired two-tailed Student’s *t*-test for B, C, D’, E’ and unpaired one-tailed Student’s *t*-test for E’’. * p < 0.05, ** p < 0.01, *** p < 0.001. ns, non-significant. PHBC, primordial hindbrain channel; LDA, lateral dorsal aorta; AAs, aortic arches; PHS, primary head sinus; CtAs, central arteries; SAG, statoacoustic ganglion; EC, endothelial cells; sib, siblings.

Since imaging revealed directed filopodial contacts from cranial vessels to sensory neurons and vice-versa (Figure 2A), we wondered about possible growth interdependency. We imaged neuronal filopodia dynamics in the absence of vessels and, endothelial filopodia in the absence of neurons. In *clo* mutants, the number of neuronal filopodia was drastically reduced (Figures 3D and 3D’). In contrast, in *neurog1* mutant embryos the number of ECs filopodia did not change, though they were shorter (Figures 3E and 3E’’). These data indicate that signals from ECs are required for neuronal filopodia formation.

While sensory neurons in SAG do not appear to be required for vascular development, our data indicate that vasculature is required for SAG neurons to proliferate during a temporal window that coincides with the period of filopodial contacts. Moreover, neuroblasts only extend filopodia when ECs are present. Altogether, the results support the hypothesis that endothelial-neuronal filopodia contacts are required for vessels to signal to neuroblasts in the SAG to restrict growth.

### Cell signalling mediated by cytonemes regulates the number of SAG neuroblasts

Cell signalling through filopodia has emerged as a novel mechanism of signalling between adjacent cells in different embryonic tissues (Kornberg, 2017; Stanganello and Scholpp, 2016; Stanganello et al., 2015). Signalling filopodia have been named cytonemes (Ramírez-Weber and Kornberg, 1999). In order to assess the possible role of filopodial signalling in SAG growth, filopodia formation was inhibited with LatrunculinB (LatB), a toxin that binds to actin and has been used to impair filopodia function *in vivo* (Morton et al., 2000; Phng et al., 2013). Imaging and quantification of filopodia in embryos treated with LatB for 6 hours indicated that the treatment was effective, the number of filopodia in ECs and in neuroblasts was reduced (Figures 4A and 4B). Interestingly, inhibition of filopodia caused an increased number of Neurod^+^ cells in LatB-treated embryos (Figure 4C) and this phenotype was also due to a higher number of Neurod^+^ cells incorporating BrdU (Figures 4D and 4E). No additional effects were found in treated *clo* mutant embryos, indicating specific effects of neuroblasts (Figure 4C). Hence, filopodia formation inhibition at early stages recapitulated the SAG growth phenotype found previously in the avascular mutant. These data demonstrate that the endothelial-neuronal filopodial contacts are cytonemes exerting a signalling role onto SAG neuroblast proliferation.

**Figure 4.**
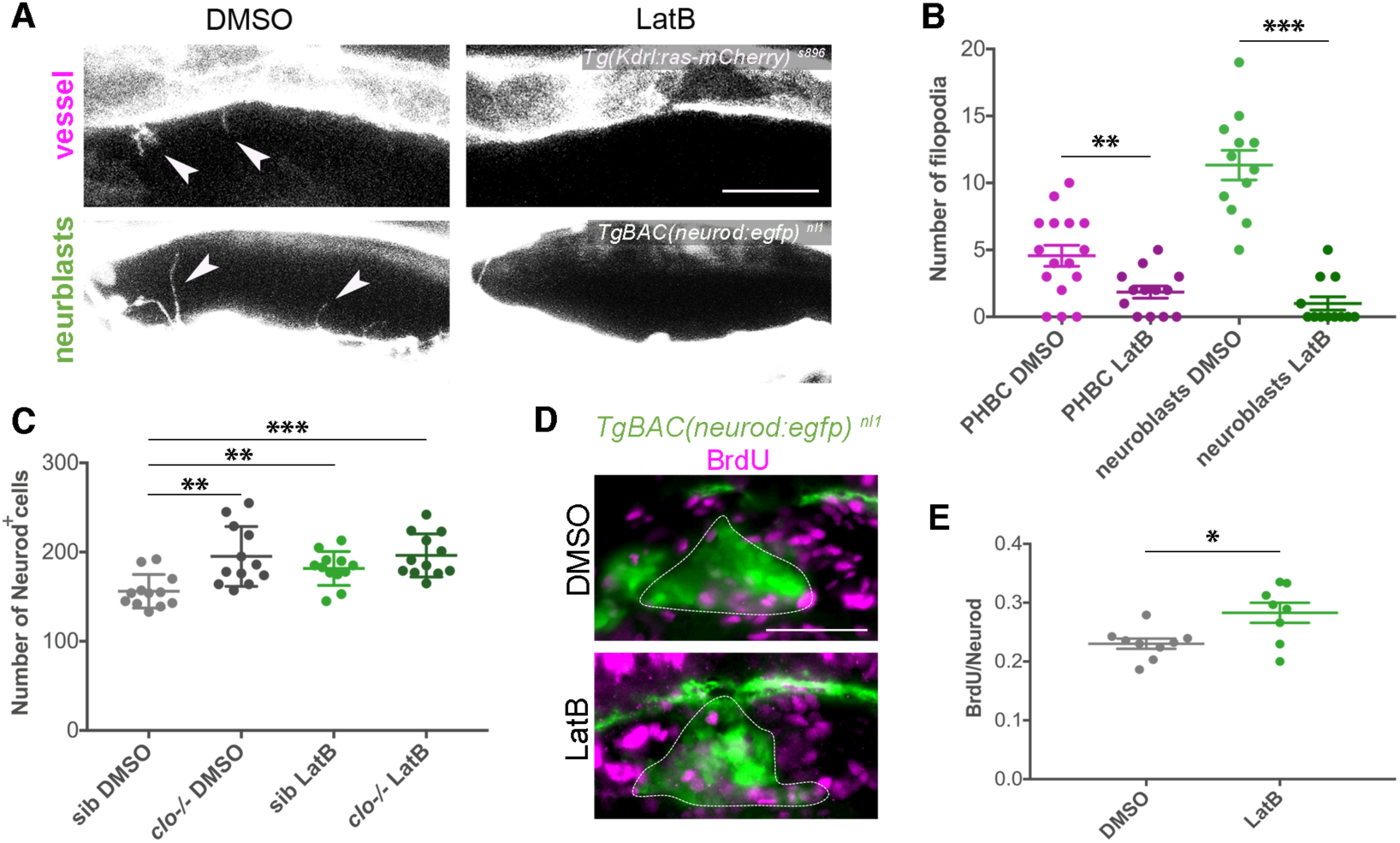
Inhibition of cytoneme processes by Latrunculin B results in expansion of neuroblasts. (A) Confocal lateral images showing filopodial processes in vessels and neuroblasts visualized through *TgBAC(neurod:egfp)* ^*nl1*^ and *(Kdrl:ras-mCherry)* ^*s896*^ respectively, after 6h treatment in DMSO or LatB, at 30 hpf. Arrowheads indicate filopodia. (B) Quantification of filopodial processes number in vessels (n = 13-16) and neuroblasts (n = 12) after 6 hours incubation with LatB compared with control (DMSO), at 30 hpf. (C) Graph showing the number of Neurod^+^ nuclei at 36 hpf, using the *TgBAC(neurod:egfp)* ^*nl1*^ background and DAPI staining, in sib and clo ^*-/-*^ embryos treated with DMSO or LatB from 24-30 hpf and washed until 36 hpf (n = 12). (D) Lateral images of BrdU incorporation (magenta) in Neurod^+^ cells of the SAG (dotted line), visualized through *TgBAC(neurod:egfp)* ^*nl1*^, in 6 hours DMSO and LatB treated embryos, at 30 hpf. (E) Graph showing the ratio of BrdU^+^ cells in the Neurod^+^ population after 6 hours DMSO (n = 9) and LatB (n = 8) treated embryos, at 30 hpf. Scale bars, 20 µm in A and 50 µm in D. Error bars, mean ± SEM. In B,E unpaired two-tailed Student’s *t*-test was performed. In C the data was analysed using one-way ANOVA, followed by post-hoc test. * p < 0.05, ** p < 0.01, *** p < 0.001. PHBC, primordial hindbrain channel; sib, siblings.

### Dll4 inhibition increases the Neurod^+^ population

To identify signals that could mediate the neurovascular crosstalk we revisited the published vascular signals regulating CNS neurogenesis for evidence of ligand expression in cranial vasculature and their receptors in cranial sensory neurons (Howe et al., 2013). In the Ventricular-Subventricular Zone (V-SVZ) of the adult brain, endothelial Jagged1 ligand restrains NSC proliferation (Ottone et al., 2014). Cranial vessels do not express Jagged1 but do express Delta-like 4 (Dll4) (Fujita et al., 2011), another Notch ligand, which is essential for proper vascular remodelling and sprouting (Bussmann et al., 2011; Leslie et al., 2007; Siekmann and Lawson, 2007). We therefore hypothesized that Dll4 could be signalling to SAG neurons, which express Notch1 at the same early timepoints (Nikolaou et al., 2009). Moreover, it has been postulated that pigmentation patterning in zebrafish is regulated by DeltaC-Notch signalling through filopodia (Hamada et al., 2014).

To test this hypothesis, we inhibited *dll4* through the well-studied *dll4* morpholino (MO) (Bussmann et al., 2011; Siekmann and Lawson, 2007) and assessed its effects on SAG growth. Remarkably, *dll4* morphant embryos presented an expansion of the Neurod^+^ population similar to the one observed previously in *clo* mutant embryos (Figure 5A). This expansion was reflected both in SAG volume (Fig 5B) as well as in Neurod^+^ cell number (Figure 5C). As a further control, *dll4* morpholino was also injected in *clo* mutant embryos and the number of Neurod^+^ cells did not change, suggesting that the results are due to loss of *dll4* expressed by the vasculature (Figures 5A-5C). These findings reveal that Dll4 is required to regulate Neurod^+^ cell number.

**Figure 5.**
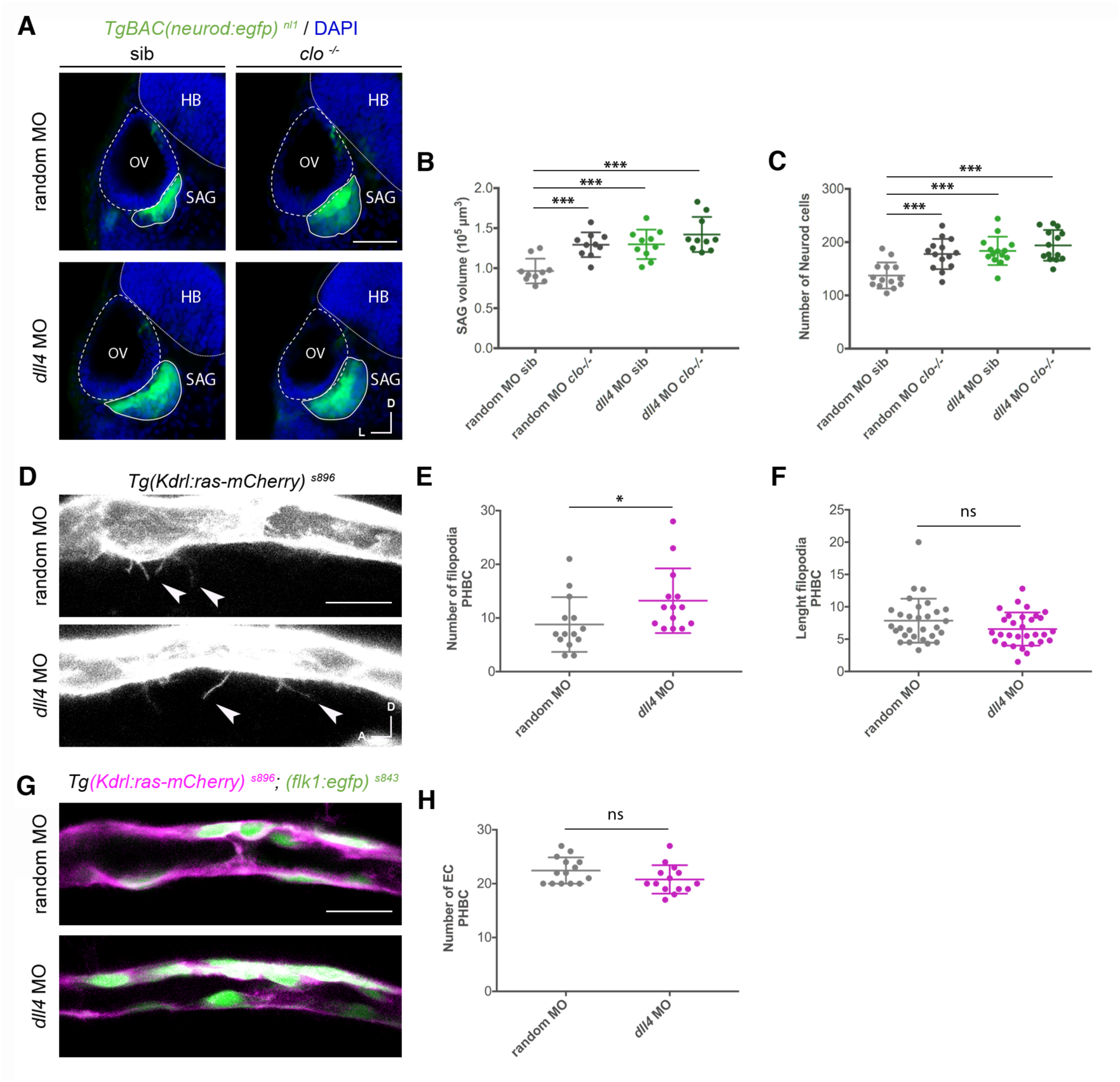
Down-regulation of Dll4 signalling induces neuroblast expansion. (A) Representative transverse sections showing the SAG in sib and clo^-/-^ *TgBAC(neurod:egfp)*^*nl1*^ embryos injected with of random or *dll4* splicing morpholino. Neurod^+^ cells: green, neuroblasts; DAPI: blue, nuclei. (B) Graph showing SAG volume quantification in sib and clo^-/-^ embryos injected with random or *dll4* splicing morpholino (n = 10), at 36 hpf. (C) Graph showing quantification of Neurod^+^ nuclei in sib and clo^-/-^ embryos injected with random or *dll4* splicing morpholino (n = 14), at 36 hpf. (D) Confocal lateral images of PHBC filopodial processes in random and *dll4* morphants visualized with the *Tg(Kdrl:ras-mCherry)* ^*s896*^, at 30 hpf. Arrowheads indicate filopodia. (E) (E,F) Quantification of PHBC filopodia number (n = 14) (E) and length (n = 14) (F), in random and *dll4* morphants, at 30 hpf. (G) Confocal lateral images showing EC membranes and cytoplasm using the *Tg(Kdrl:ras-mCherry)* ^*s896*^ and *Tg(flk1:egfp)* ^*s843*^ respectively, in the PHBC, at 30 hpf. (F) Quantification of EC number in random and *dll4* morphants (n = 14), at 30 hpf. Scale bars, 50 µm in A, and 20 µm in D,G. Error bars, mean ± SEM. In B,C the data was analysed using one-way ANOVA, followed by post-hoc test. In E the non-parametric Mann-Whitney *U* test was performed. In F,H unpaired two-tailed Student’s *t*-test was used. * p < 0.05, ** p < 0.01, *** p < 0.001. HB, hindbrain; OV, otic vesicle; SAG, statoacoustic ganglion; PHBC, primordial hindbrain channel; EC, endothelial cell.

In angiogenic sprouts, the number of filopodia increases after *dll4* downregulation (Suchting et al., 2007). When we assessed the number of filopodia in the developed PHBC, we detected a small increase in filopodial number and no differences in their length (Figures 5D-5F). Thus, the increase of Neurod^+^ cells in *dll4* morphants cannot be attributed to a lack of filopodial processes but to a specific lack of signal. Finally, the number of ECs did not change in *dll4* morphant embryos compared to embryos injected with a random morpholino (Figures 5G and 5H). Therefore, the increase in Neurod^+^ cell number after *dll4* down-regulation is not due to vascular developmental changes but to Dll4-Notch1 signalling.

Together, these results suggest that Dll4 signalling constitutes one of the key molecular mechanisms by which the vasculature regulates cranial sensory neurogenesis.

### Absence of vasculature impairs sensory neuronal differentiation

Since more neuroblasts form in the SAG in the absence of vasculature, we next assessed if these neuroblasts differentiate, resulting in a higher number of Islet2^+^ post-mitotic cells at later stages. Unexpectedly we found, after plotting the number of immunostained Islet2^+^ nuclei from 30 to 96 hpf, that there was a significantly lower number of Islet2^+^ cells in *clo* mutants from 54 hpf onwards (Figures 6A and 6B). The differentiation curves indicate that, while at early timepoints sensory differentiation occurs partially independently of vasculature, from 54 hpf onwards, cranial vasculature is required to promote differentiation of Neurod^+^ cells to Islet2^+^ cells. As expected from the immunostaining data, the fluorescence from sensory axons was also reduced in *clo* mutant embryos, labelled by the *Tg(isl2:GFP)* ^*zc7*^ reporter line which marks *in vivo* the cytoplasm of differentiated sensory neurons (Pittman et al., 2008) (Figure 6C).

**Figure 6.**
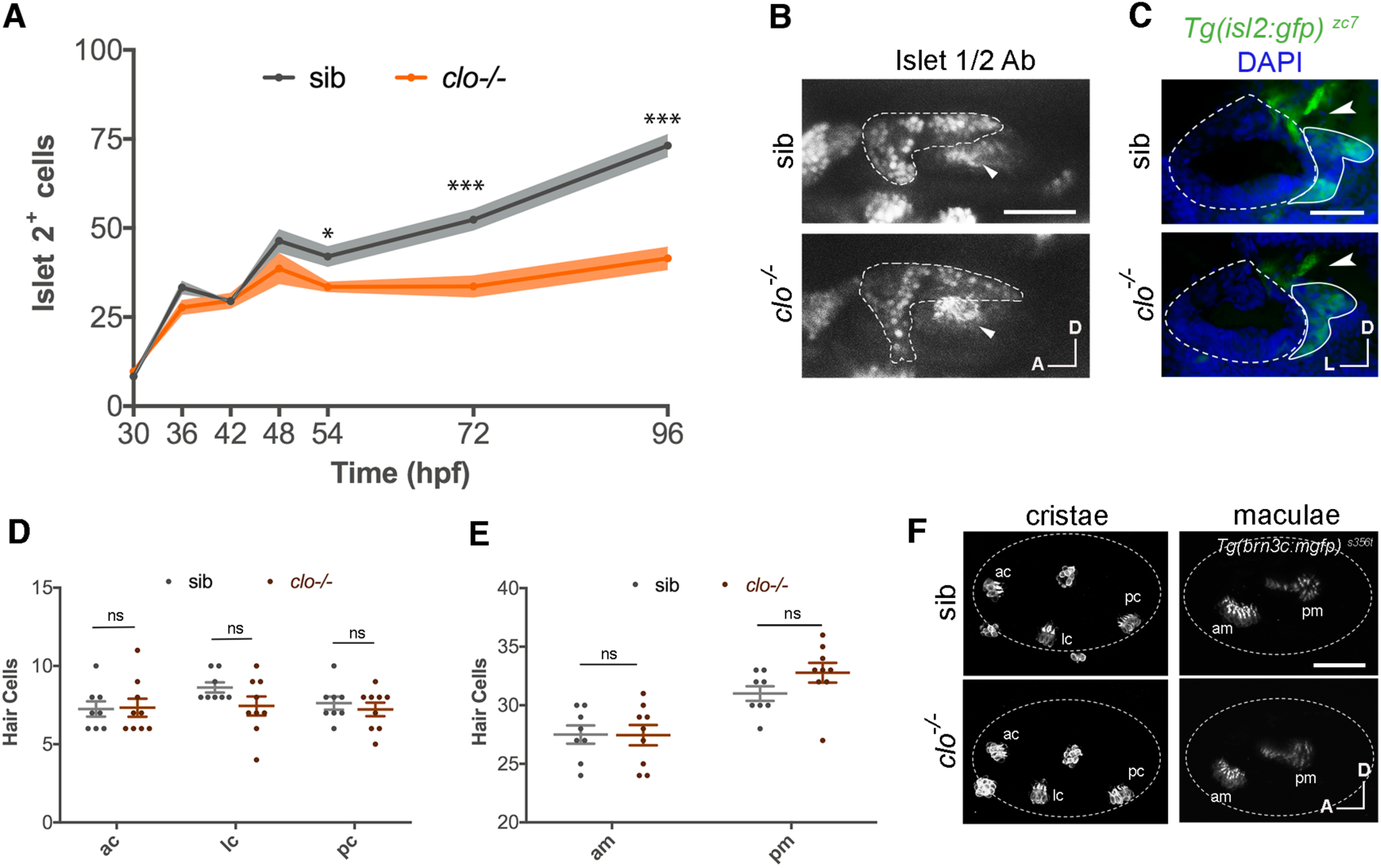
Sensory neuronal differentiation requires blood vessels. (A) Graph showing the number of Islet 2^**+**^ nuclei in the SAG at different timepoints from 30 to 96 hpf (n = 4-10). (B) Confocal lateral images of immunostained Islet2^**+**^ nuclei in the SAG (dotted line, visualized by *TgBAC(neurod:egfp)* ^*nl1*^) in sib and clo^**-/-**^ embryos at 72 hpf. Arrowheads indicate pm. (C) Transverse sections showing axon density (arrowhead) in the *Tg(isl2:gfp)* ^*zc7*^ in sib and clo^**-/-**^ embryos, of cryostat sections, at 72 hpf. Dotted line indicates OV and continuous line the SAG. (D,E) Graph showing quantification of hair cell number in cristae (D) and maculae (E) in sib (n = 8) and clo^**-/-**^ (n = 9) embryos. (F) Representative lateral images of hair cells in cristae and maculae using the *Tg(brn3c:mGFP)* ^*s356t*^, at 72 hpf. Dotted line indicates otic vesicle. Non-labelled hair cells are neuromasts. Scale bars, 50 µm. Error bars, mean ± SD for A; and mean ± SEM for D,E. Unpaired two-tailed Student’s *t*-test. * p < 0.05, ** p < 0.01, *** p < 0.001. ns, non-significant. OV, otic vesicle; SAG, statoacoustic ganglion; ac, anterior crista; lc, lateral crista; pc, posterior crista; am, anterior macula; pm, posterior macula; sib, siblings.

To determine if cranial vasculature regulates neurogenesis in other sensory ganglia besides the SAG, we assessed for filopodia contacts and quantified the number of Neurod^+^ and Islet2^+^ cells in different ganglia. Here, we did not perform a temporal analysis, but we chose discrete timepoints. At 30 hpf, we could also detect filopodia from the PHBC to the trigeminal and anterior lateral line ganglia (Figure S4C). At 36 hpf, we found a significantly increased number of Neurod^+^ cells in the trigeminal ganglion of *clo* mutants (Figures S4A and S4B), whereas at 72 hpf there was a reduced number of Islet2^+^ cells in the trigeminal and vagal ganglia (Figures S3D-S3G). Therefore, the neurovascular interaction appears to extend to other sensory ganglia.

To complete the analysis, we assessed if vasculature could regulate the development of other cell types such as the hair cells of the inner ear. For this aim, we generated *clo* homozygous mutants in the *Tg(brn3c:mGFP)* ^*s356t*^ background that labels the membranes of hair cells and their kinocilia (Xiao et al., 2005). Serial confocal stacks taken at 72 hpf and quantification of hair cells did not reveal significant differences in hair cell number in cristae nor maculae of *clo* mutants compared to siblings (Figures 6D-6F). Therefore, vasculature is specifically affecting neuronal differentiation of the inner ear but not hair cell number.

### Initiation of blood flow and oxygen sensing triggers sensory differentiation

It has been reported that in zebrafish blood flow begins due to the completion of anastomosis of cranial blood vessels around 48 to 60 hpf (Siekmann et al., 2009). The coincidence of this timing with the one of sensory differentiation defects in *clo* mutant embryos prompted us to explore whether the inducing signal of sensory differentiation was linked to blood flow. To this aim, we incubated embryos with Nifedipine, a drug that inhibits heartbeat and blood flow (Li et al., 2008). As expected from blood flow inhibition, we observed a drastic vessel diameter reduction in Nifedipine-treated embryos (Figures 7A and 7B). In embryos treated with Nifedipine from 54 to 72 hpf, the number of Islet2^+^ cells decreased, recapitulating the defects found in *clo* mutant embryos (Figures 7C-7E). Treatment in the avascular model did not further enhance the phenotype, suggesting that the differentiation arrest is mostly attributed to the absence of blood flow in *clo* mutant embryos and not associated to side effects of Nifepidine (Figure 7E). The same results were obtained when treatment was done with BDM, another drug used to inhibit blood flow (Anderson et al., 2008) (Figure S5C and S5D). To narrow down the time window of blood flow requirement, we incubated embryos with Nifedipine from 60 to 72 hpf and from 48 to 60 hpf. Incubation from 60 to 72 hpf did not inhibit differentiation (Figures 7F and 7G), whereas incubation from 48 to 60 hpf did (Figures S5A and S5B). These results indicate that blood flow induction of sensory differentiation takes place in a narrow time window and once signalling is initiated, blood flow is no longer required.

**Figure 7.**
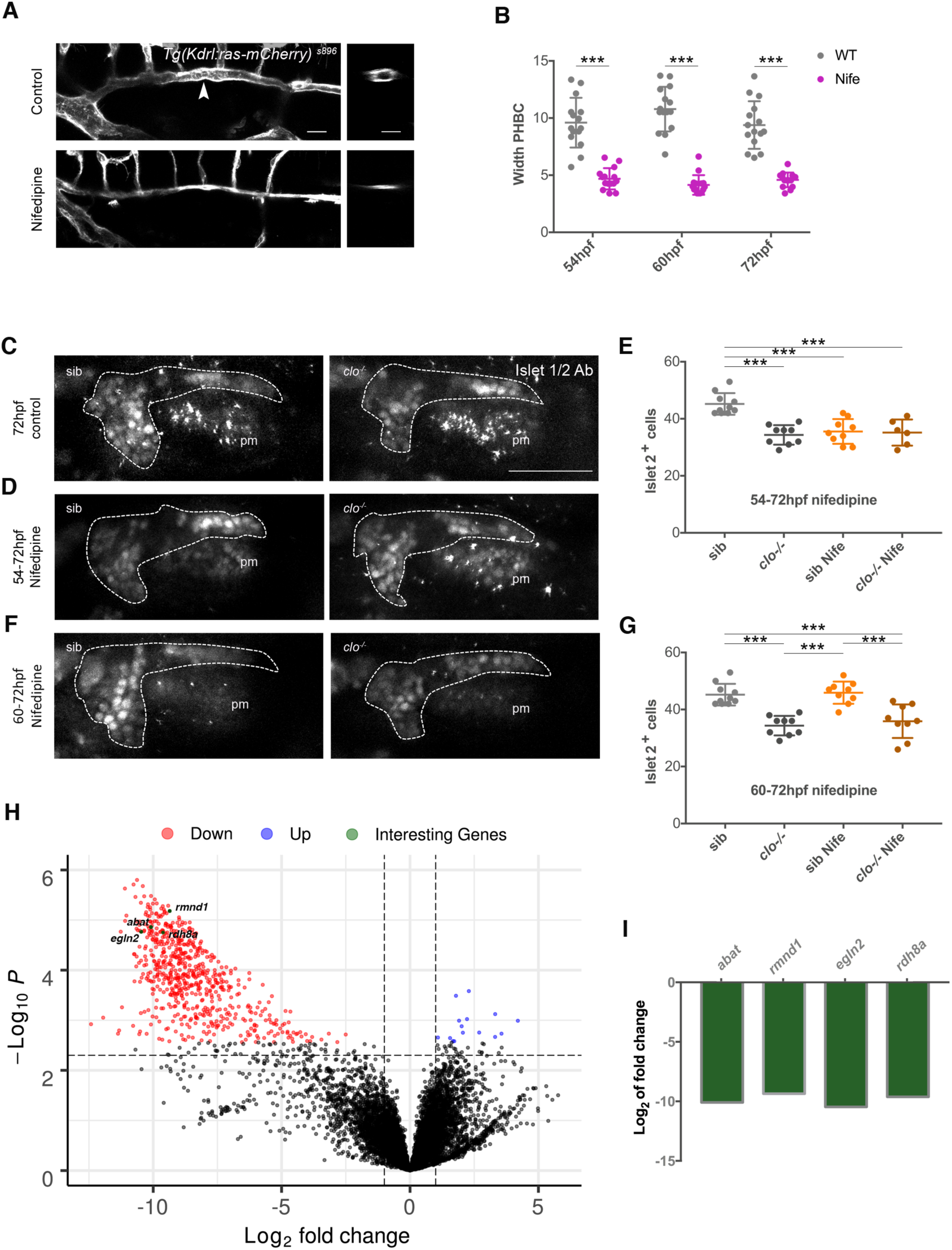
Blood flow triggers transcriptional changes related to oxygen sensing and metabolism together with sensory neuronal differentiation in a restricted temporal window. (A) Representative images of the PHBC of control and 48-54 hpf Nifedipine treated *Tg(Kdrl:ras-mCherry)* ^*s896*^ embryos, at 54 hpf, in lateral (right) and transversal optic reconstructions (left) in the point marked by the arrowhead. (B) Analysis of PHBC width in control and Nifedipine treated *Tg(Kdrl:ras-mCherry)* ^*s896*^ embryos, at 54, 60 and 72 hpf with treatment starting at 48 hpf (n=15). (C-G) Confocal lateral images showing immunostained Islet2^**+**^ nuclei in the SAG (dotted line, visualized by *Tg(neurod:egfp)* ^*nl1*^) in sib and clo^**-/-**^ embryos at 72 hpf in control (untreated) (C), 54-72 hpf Nifedipine treated embryos (D), 60-72 hpf Nifedipine treated embryos (F), and quantified in E,G. 72 hpf sib (n = 10), 72 hpf clo^**-/-**^ (n = 9), 54-72 hpf sib Nife (n = 9), 54-72 clo^**-/-**^ Nife (n = 6), 60-72 hpf sib Nife (n = 9), 60-72 hpf clo^**-/-**^ Nife (n = 9). (H) Volcano plot of differentially expressed genes (DEGs) in the SAG at 54 hpf. Red and blue plots represent significant downregulated and upregulated DEGs, respectively (P<0.05; FC<-1; FC>1), while black plots represented insignificant DEGs (P≥0.05; FC≥-1; FC≤1). Genes of special interest are highlighted in green. (I) Histogram showing the fold change of DEGs of special interest in clo ^*-/-*^ versus sibling SAGs at 54 hpf. Scale bars, 50 µm for A,B,D,F, and 20 µm for H. Error bars, mean ± SEM. Unpaired two-tailed Student’s *t*-test. * p < 0.05, ** p < 0.01, *** p < 0.001. ns, non-significant. pm, posterior macula; Nife, Nifedipine; sib, siblings; PHBC, primordial hindbrain channel.

To obtain insight into the molecular mechanisms underlying the arrest in sensory neuron differentiation in the absence of vasculature, we conducted a transcriptional profiling of freshly sorted Neurod GFP^+^ SAG cells from *clo* mutant and sibling embryos at 54 hpf. RNA-seq analysis revealed a transcriptional profile response in the absence of vasculature with 641 genes significantly downregulated and 15 upregulated in clo mutant compared to control embryos (Figure 7H). This suggest that in absence of blood flow some signalling pathways are shut down rather a new transcriptional program being activated, what matches with a cell differentiation arrest. Whereas we expected pathways related to differentiation signals or cell-cycle control being down-regulated, most differentially expressed genes found where genes related with oxygen sensing (egln2), reduction-oxidation (redox) process (rdh8a) and mitochondria metabolism (rmnd1 and abat) (Figures 7H ans 7I). Thus, the first transcriptional response to blood flow onset seems to be related to oxygen sensing and switch from glycolysis to OxPhos that ultimately drives sensory neuronal differentiation, as observed in other contexts (Hawkins et al., 2016; Homem et al., 2014; Kida et al., 2015; Lange et al., 2016; Ryall et al., 2015).

Altogether, the results define for the first time a prominent role of cranial vasculature in sensory neurogenesis that can be segregated into two sequential and independent mechanisms. A first one, in which Dll4-Notch1 cytoneme signalling from ECs to neurons regulates neuroblast quiescence, and a second one in which initiation of blood flow and oxygen sensing induces a metabolic switch and sensory neuron differentiation.

## Discussion

The SAG and other cranial sensory ganglia reside in a vascularized milieu in embryos and adults. A fundamental question in developmental neurobiology is whether or not vasculature, in addition to transport of nutrients and oxygen, provides instructive cues for sensory neurogenesis. This is relevant to understand hearing loss and other sensory neuropathies linked to vascular defects (Jensen et al., 2004; Lago et al., 2018; Paul et al., 2016). Our detailed temporal analysis reveals that cranial ganglia indeed require vascular signals for proper growth and differentiation. We identified two segregated mechanisms: an early cytoneme-based, cell-cell contact dependent mechanism regulating neuroblast proliferation and a later blood flow-dependent mechanism controlling neuronal differentiation.

In the V-SVZ, quiescent NSCs physically contact blood vessels with specialized endfeet (Kacem et al., 1998). By taking advantage of the imaging and cell reporter tools available in the zebrafish model we resolve, for the first time, that cell-cell communication between vasculature and neuroblasts takes place through dynamic cytoneme extensions that project between them. It is still possible that in other contexts, dynamic contacts also exist, but imaging in fixed tissue might have hampered their identification, as suggested by Obernier and colleagues (Obernier et al., 2018). Filopodia in angiogenic tip cells and migrating ECs have been described (Fantin et al., 2015; Gerhardt et al., 2003; Ochsenbein et al., 2016; Phng et al., 2013). To our knowledge, filopodia from endothelial stalk cells contacting neuronal cells have not been reported before. One could envision cell-cell contact with only ECs extending filopodia to neuronal somas but, in our system, both cell types project directional filopodia. Signalling filopodia or cytonemes in *Drosophila* and vertebrates have been demonstrated, extending the cell-cell signalling distance of influence (Kornberg, 2017; Stanganello and Scholpp, 2016; Stanganello et al., 2015). This is relevant in the facial mesenchyme where vasculature does not grow inside the sensory ganglia but adjacent. Different types of cytoneme contacts have been described in previous studies, in some cases cytonemes contact the cell body of their target cell (Huang and Kornberg, 2015; Stanganello et al., 2015), while in other cases cytoneme-cytoneme contacts are observed (Du et al., 2016; González-Méndez et al., 2017; Huang and Kornberg, 2015; Roy et al., 2014; Sanders et al., 2013). It is difficult to tell if in our system cytoneme-cell body contacts occur, but we have clearly observed cytoneme-cytoneme contacts. These cytonemes have distinct behaviours, either highly transient or more stable, the latter possibly being actively signalling.

Moreover, the study highlights the requirement of signals from ECs for neuronal cytoneme growth. By blocking filopodia projections, we demonstrate that they have a signalling function in the negative control of neuroblast cell proliferation. Most likely only a subset of neuroblasts are contacted by endothelial cytonemes, allowing a scattered pattern of cell cycle-arrested neuroblasts between others with proliferative capacities. Cytonemes have been described to participate in stem cell maintenance in *Drosophila* (Fuwa et al., 2015; Inaba et al., 2015; Mandal et al., 2007; Rojas-Ríos et al., 2012), but not yet described to participate in a NSC niche in vertebrates.

In contrast to other contexts in which neurons control vascular patterning (Himmels et al., 2017), our results suggest that in the context of sensory ganglia and surrounding vessels, an absence of cranial ganglia does not have a major effect on early vasculogenesis.

During adult neurogenesis, cell contact-dependent neurovascular signalling is mediated by ephrinB2 and Jagged1 present in ECs that suppress cell-cycle entry in NSCs (Ottone et al., 2014). We propose that in the PNS Dll4, but not Jagged1, plays a similar role. Maintenance of a quiescent and undifferentiated population of neural progenitors is key for life-long regeneration. Thus, the role we have uncovered for cranial vasculature and Dll4 in the cranial sensory niche might be relevant in regenerative studies upon neuronal damage.

Several studies have highlighted the role of blood flow initiation in remodelling the vasculature, VEGF expression, angiogenesis and stimulation of hematopoietic stem cell formation (Adamo et al., 2009; Bussmann et al., 2011; Nicoli et al., 2010). Blood flow can exert tangential and perpendicular forces on ECs leading to actomyosin rearrangements, YAP-TAZ nuclear translocation and the induction of the mechano-sensitive transcription factor Klf2a (Nakajima et al., 2017; Nicoli et al., 2010; Le Noble et al., 2008). Ultimately, blood flow triggers gene transcriptional changes in ECs and vessel maturation (Chiu and Chien, 2011).

On the other hand, blood vessels can exert their roles on neural cells via delivery of nutrients or oxygen (Cleaver and Dor, 2012; Ramasamy et al., 2015). The transcriptome analysis of SAG neuroblasts in clo mutant vs control embryos at the timepoint of blood flow initiation allowed us to identify *egln2* (*Egl nine homolog 2*) as one of the early-response genes more differentially expressed. Egln2 is cellular oxygen sensor involved in the degradation of the transcription *factor hypoxia-inducible factor* (*HIF1*) (Epstein et al., 2001), thus in wild-type conditions embryos sense the passage from hypoxic to normoxic conditions. Additionally, genes related to redox state and mitochondria metabolism are also differentially expressed. Our data suggest that SAG sensory precursors switch from glycolytic to OxPhos to induce their differentiation. Numerous populations of multipotent SC, i.e hemaepoietic SC, intestinal crypt SC and neural SC, use a glycolytic program in their niche and switch to OxPhos upon regeneration and differentiation (Cliff and Dalton, 2017; Lange et al., 2016; Mohrin et al., 2015; Shi et al., 2015).

Our current work identifies novel mechanisms for head vasculature in the balance between sensory precursor and differentiation fates and shall contribute to our basic understanding of the cranial sensory niche in which head vasculature enters in scene.

## Acknowledgments

We thank D Stainer for the *Tg(cloche)* ^*m378*^ and *Tg(kdrl:ras-mCherry)* ^*s896*^ lines, the PRBB Imaging Facility and the UPF Genomics Core and Flow Cytometry Facilities for technical support, T. Schilling, A. Bigas and I. Fariñas for comments on the manuscript and lab members for discussions. The work was supported by BFU2014-53203 and BFU2017-82723P from MCINN to BA and the Unidad de Excelencia Maria de Maeztu (MDM-2014-0370).

## Author Contributions

LT and BA designed research, LT and AB performed experiments, LT, AB and BA analysed data and TL and BA wrote the manuscript.

## STAR Methods

### Zebrafish Strains and Genotyping

Zebrafish (*Danio rerio*) were maintained as previously described (Westerfield, 2000) at the aquatic facility of the Parc de Recerca Biomèdica de Barcelona (PRBB). The following transgenic lines in different combinations were used: *Tg(neurog1)* ^*hi1059*^ (Golling et al., 2002) mutant to inhibit neurogenesis and *Tg(cloche)* ^*m378*^ (Stainier et al., 1996) as an avascular model. Five different reporter lines were used: *Tg(Kdrl:ras-mCherry)* ^*s896*^ (Chi et al., 2008) and *Tg(flk1:EGFP)* ^*s843*^ (Jin et al., 2005) for vessels, *TgBAC(neurod:EGFP)* ^*nl1*^ (Obholzer et al., 2008) for neuroblasts, *Tg(isl2:gfp)* ^*zc7*^ (also called Isl3) (Pittman et al., 2008) for sensory differentiated neurons and *Tg(brn3c:mGFP)* ^*s356t*^ (Xiao et al., 2005) for hair cells.

Embryos were staged as previously described (Kimmel et al., 1995). Adult heterozygous for *neurog1* ^*hi1059*^ allele were identified by PCR genotyping by fin-clip genomic DNA(Golling et al., 2002). Homozygous *neurog1* mutant embryos were recognized by the significant reduction of GFP signal in *Tg(neurog1)* ^*hi1059*^, *BAC(neurod:EGFP)* ^*nl1*^, *(Kdrl:ras-mCherry)* ^*s896*^ background. Avascular mutants were identified by the lack of vasculature in the reporter line for endothelial cells *Tg(Kdrl:ras-mCherry)* ^*s896*^ or *Tg(flk-1:EGFP)* ^*s843*^, lack of blood cells and circulation (visible under scope from 30-36hpf) and atrium enlargement, as described by Stainer et al. (Stainier et al., 1995).

In both cases, wild-type and heterozygous-deficient siblings were undistinguishable and both have been used and referred as siblings (sib). Finally, since *clo* mutant embryos die around 7dpf (Stainier et al., 1996), we studied embryos until 4 dpf to exclude vessel directed effects from embryonic morbidity.

### Live imaging, confocal microscopy and quantifications

Images of cryosections were acquired with a Leica DFC 7000 T microscope, and 40x oil immersion (0.75 NA).

Whole-mount embryos imaging was performed on a Leica SP8 confocal microscope, with 20x glycerol immersion lens (0.7 NA). Embryos were mounted in 1% low-melting agarose (Ecogene) in PBS or Danieau’s solution, for fixed and life embryos respectively, and placed onto a glass-bottom dish. For life embryos, 0.1% Tricaine (Sigma) was also added. The 488- and 561-nm laser lines were employed at a scan speed of 600 Hz.

For time-lapse imaging of filopodial, the resonant mode (8000 Hz) was used to achieve a high spatiotemporal resolution. Frames were captured simultaneously with 488 nm and 561 nm lasers at a time interval of 15 seconds. Z-size of 2 µm was used for videos and 0.5 µm for static stacks and transverse optical reconstructions. Confocal stacks and movies were flattened by maximum projections using FIJI (Image J).

Measurements were done manually with FIJI (Image J) for SAG and PHBC filopodia number, length and orientation using one single static image, from a defined region of interest (ROI). PHBC width was measured from maximum projection confocal images always at the same point, between CtA 2 and 3. PHBC EC number was calculated by perinuclear green cytosolic signalling from *Tg(flk-1:EGFP)* ^*s843*^ and EC contours from *Tg(Kdrl:ras-mCherry)* ^*s896*^ in a defined ROI. When ROIs were used, the same area was used to compare samples. SAG anteroposterior (AP) and dorsoventral (DV) axes length were measured from maximum projection confocal lateral images. The mediolateral (ML) SAG axis was calculated from 5 serial cryosections, in which the biggest measurement was used.

The number of Neurod^+^ nuclei was counted manually using DAPI staining in the *TgBAC(neurod:egfp)* ^*nl1*^ labelled cells. *neurog1*^*+*^ cells were also counted manually after an ISH and DAPI staining. The number of HC was calculated using the *Tg(brn3c:mGFP)* ^*s356t*^ and counting kinocilia number.

The volume of the SAG was measured by calculating the area the 5 serial cryosections that include all SAG and multiplying them with each section thickness (20µm).

### Statistical Analysis

All the data in the work were first tested for normal distribution using the Kolmogorov-Smirnov test and the Levene’s test for homogeneity of variances. For two-group comparison two-tailed Student’s *t*-test was used or Mann-Whitney *U* test for non-parametric tests. When more than two samples were being compared, one-way ANOVA followed by post-hoc test was performed. Values are expressed as median ± SEM. Graphs were performed with PRISM 7 (Graph-Pad) software. Error bars represent SEM, but for Fig. 2d and Fig. 5a, were shaded error bars represent SD. ns= non-significant, *p≤0.05, **p≤0.01, ***p≤0.001.

### Immunostaining and *in situ* Hybridization

Embryos at the desired stage were fixed with 4% paraformaldehyde (PFA) in PBS 0,1% Tween-20 (PBT) for 3 hours at RT or overnight at 4°C. Fixed embryos were dehydrated, kept in 100% MeOH −20°C 1h or overnight (O/N), and rehydrated. Embryos were permeabilized with Proteinase K 1:1000 in PBT, or cold Acetone for mouse anti-Islet1/2, and incubated with blocking solution (BS) for 2h at room temperature (RT) and with primary antibody O/N at 4°C. Primary antibody was washed 5 times 10min, and secondary antibody was applied for 2h RT or O/N 4°C.

Embryos were cryoprotected in 15% sucrose and embedded in 7.5% gelatine/15% sucrose. Blocks were frozen in 2-methylbutane (Sigma) to improve tissue preservation. Sagittal and transversal sections were cut at 20µm thickness in a Leica CM 1950 cryostat.

For BrdU incorporation experiments, dechorionated live embryos were incubated with 10mM BrdU, 10% DMSO in Danieau’s solution for 6 hours. After this they were washed twice in clean Danieau’s solution for 5 min prior 4% PFA fixation for 3 hours at RT. BrdU staining was performed in cryosections after 2N HCl DNA denaturation 1h RT. Anti-BrdU (bdbiosciences; mouse Cat: 555627, 1:100). Anti-GFP was used to enhance GFP signal from transgenic lines (Torrey Pines BioLabs Inc; rabbit Cat: TP401, 1:400). Anti-Islet1/2 was used to detect differentiated neurons (Cat: TP401; mouse Cat: 39.4D5, 1:200). Alexa-fluor 488 and 594 anti-rabbit and anti-mouse (Invitrogen, 1:400) were used as secondary antibodies. DAPI staining for cryosections was done at 1:10000 PBS for 5min, and 2×5min wash in PBS followed by mowiol mounting.

Whole-mount *in situ* hybridization was performed as previously described (Thisse et al., 2004) using *neurog1* digoxigenin-labelled probe.

### Inhibitor treatments

To block actin polymerization in filopodia, dechorionated live embryos were incubated in 0,1µg/mL Latrunculin B (LatB) (Merk, Millipore), 0,4% DMSO in Danieau’s solution for 6 hours (Phng et al., 2013); while control embryos were raised at 0,4% DMSO only. After treatment, embryos were washed twice in clean Danieau’s solution for 5min followed by immediate confocal imaging for filopodia visualization or 4% PFA fixation. In other cases, embryos were washed in Danieau’s solution until 36 hpf. It has been previously demonstrated that the concentration used of LatB specifically inhibits filopodial formation while it does not affect other cellular behaviours for up to 17 hours of treatment (Phng et al., 2013).

Blood flow was blocked through the use of two different drugs. Dechorionated live embryos were incubated for 6, 12 and 24 hours in 40µM Nifedipine or 20mM BDM (2, 3-butanedione-2-monoxime) (Sigma-Aldrich) in Danieau’s solution, washed twice 5min and fixed in 4% PFA at the desired stage (Anderson et al., 2008; Li et al., 2008).

### Morpholino knockdown experiments

To inhibit protein translation of Dll4, 10ng of random and *dll4* splicing morpholino antisense oligonucleotides (Gene Tools, Philomath, OR) were injected in the yolk of 1-2 cell stage embryos as previously described (Siekmann and Lawson, 2007). Efficiency of the knockdown was assessed by RT-PCR (data not shown).

### FACS purification of GFP^+^ cells

Embryos of the transgenic line *Tg(neurod:gfp)* ^*nl1*^ were grown until 54 hpf, stage at which they were anesthetized with tricaine. The SAG was dissected cutting the embryos head after the eye and in the posterior end of the otic vesicle. The hindbrain was also removed to prevent the sorting of any neurod^+^ cells also present in the CNS. The presence of the SAG was confirmed under a fluorescent scope. The dissected tissue was kept in Medium 199, on ice, until we summed around 60 embryos per each condition siblings and *clo*. Cell were disaggregated using a protease solution (10x Trypsin, 0,5M EDTA, 1x PBS) for 20 min with mechanical help by both vortexing and doing up&down with a 200 µL pipete. Cell disaggregation was stopped with STOP solution (30% FBS, 2M CaCl2, 1x PBT). Cells were recovered by centrifugation and re-suspended in cell suspension medium (20% BSA, 0,5M EDTA, 1x PBS). Neurod^+^ cells were FACS purified based on GFP fluorescence utilizing a INFLUX Sorter. Dead cells were eliminated by DAPI staining (1:1000). Cells were collected in collection medium (1:1, PBS:FBS) and immediately used to RNA extraction.

Four replicates were used, always preparing one sample for siblings and another for *clo* mutants each time, coming from the same breed.

### RNA extraction and sequencing

RNA-seq was performed on FACS-sorted neuroblasts from the *Tg(neurod:egfp)* ^*nl1*^ zebrafish embryos at 54 hpf. RNA from these cells was extracted using the PicoPure(tm) RNA Isolation Kit (Thermo Fisher, KIT0214).

Cells from around 60 embryos were pooled to obtain enough RNA for library preparation. cDNA was prepared with SMART-Seq® v4 Ultra® Low Input RNA Kit for Sequencing (Takara, 634888) and validated using Bioanalyzer High Sensitivity DNA Kit (Agilent, Cat. No. 5067-4626). Between 66-306 pg of these cDNAs were used to prepare libraries using Nextera XT DNA Library Prep Kit (Illumina, FC-131-1024). Libraries were validated using Bioanalyzer High Sensitivity DNA Kit (Agilent, Cat. No. 5067-4626). Then, libraries were normalized and pooled for sequencing using the NextSeq High Output (Illumina, 20024907) with 75 bp paired ends run with 50 million expected reads per sample.

Mapped reads with the Zebrafish genome (version GRCz11) were obtained using the STAR software (v2.5.2b, https://github.com/alexdobin/STAR), and the reads’ counts within exons with the HTSeq software (v0.9.1, https://htseq.readthedocs.io/en/release_0.11.1/). The DE analysis was done with the limma package in R (v3.38.3, https://bioconductor.org/packages/release/bioc/html/limma.html).

## Supplemental Figures and Video Legends

**Figure S1.**
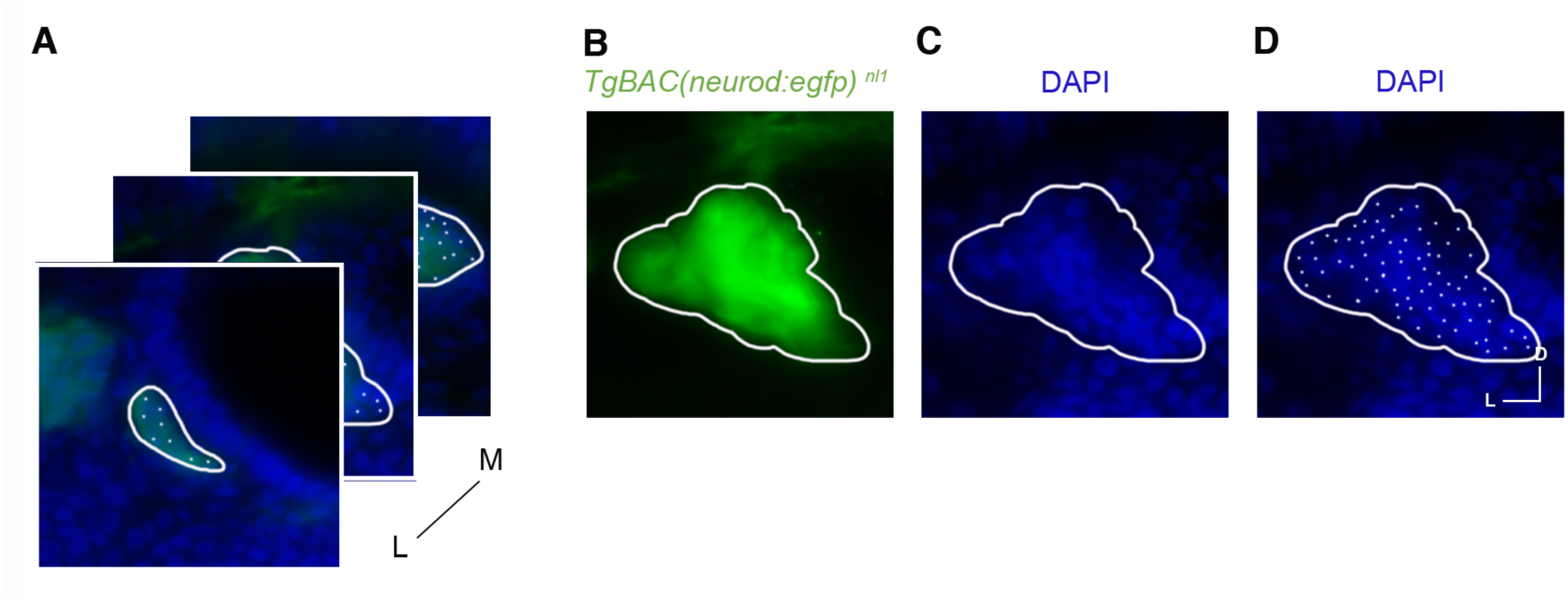
Protocol used for quantification of Neurod^+^ cell number. (A-C) For a series of latero-medial in sagittal or antero-posterior transversal (data not shown) cryostat sections, the perimeter of the SAG was drawn using the green signal of TgBAC(neurod:egfp) nl1 (B), subsequently green light was hidden and blue light from DAPI staining was visualized (C) to manually count the number of cells (D).

**Figure S2.**
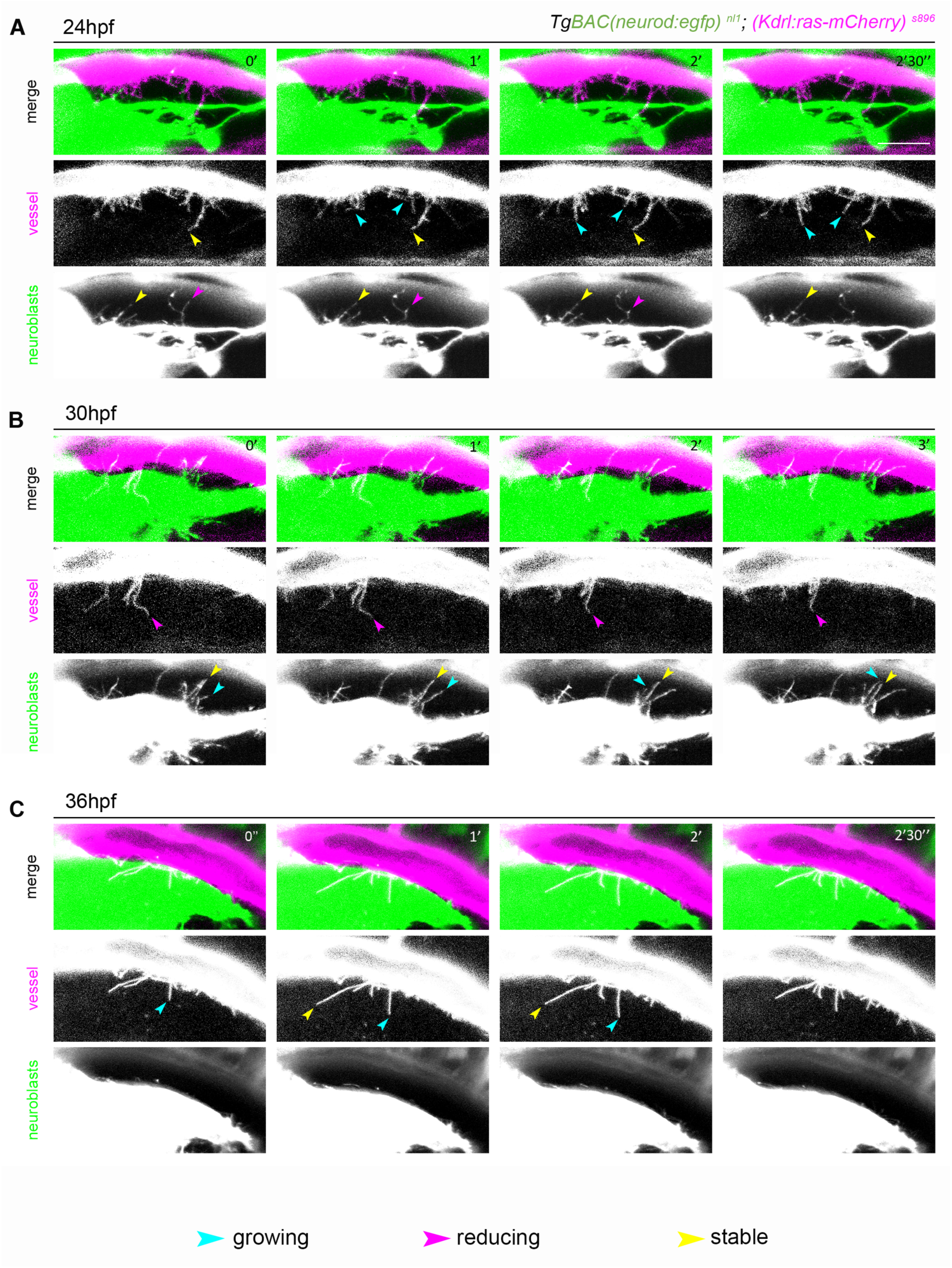
EC from the PHBC and neuroblasts from the SAG extend filopodia towards each other from 24 to 36 hpf. (A-C) Still images of time-lapse videos of *TgBAC(neurod:egfp)* ^*nl1*^; *(Kdrl:ras-mCherry)* ^s896^ showing filopodia extending from the PHBC to neuroblasts and from neuroblasts to PHBC at 24 hpf (A), 30 hpf (B), and 36 hpf (C) (time-lapses are representative of 5 experiments), in a lateral view. Cyan, magenta and yellow arrowheads point to examples of growing, reducing and stable filopodia, respectively. Scale bar 50 µm. PHBC, primordial hindbrain channel.

**Figure S3.**
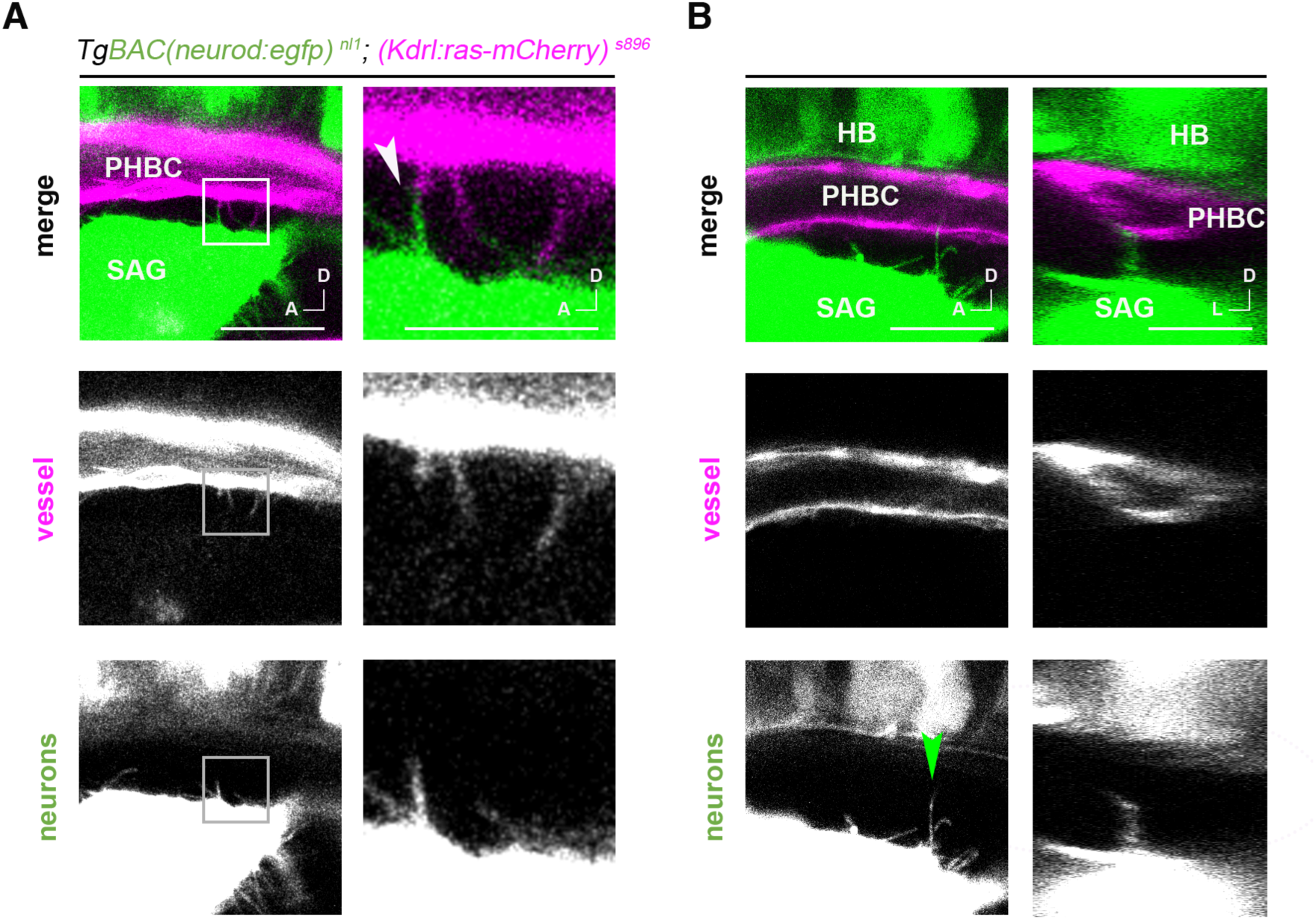
Filopodia from vasculature and neuroblasts contact each other at the same plane. (A) Confocal lateral image of a single plane showing a filopodia-filopodia contact between EC and neuroblasts. Square shows magnified image in the right, and arrowhead the mentioned contact. (B) Confocal lateral image of a single plane and transverse optical reconstruction also in single plane showing a filopodial process from the neuroblasts contacting the EC. Arrowhead indicates point for transverse optical reconstruction. Scale bar 20 µm. PHBC, primordial hindbrain channel; SAG, statoacoustic ganglion; HB, hindbrain.

**Figure S4.**
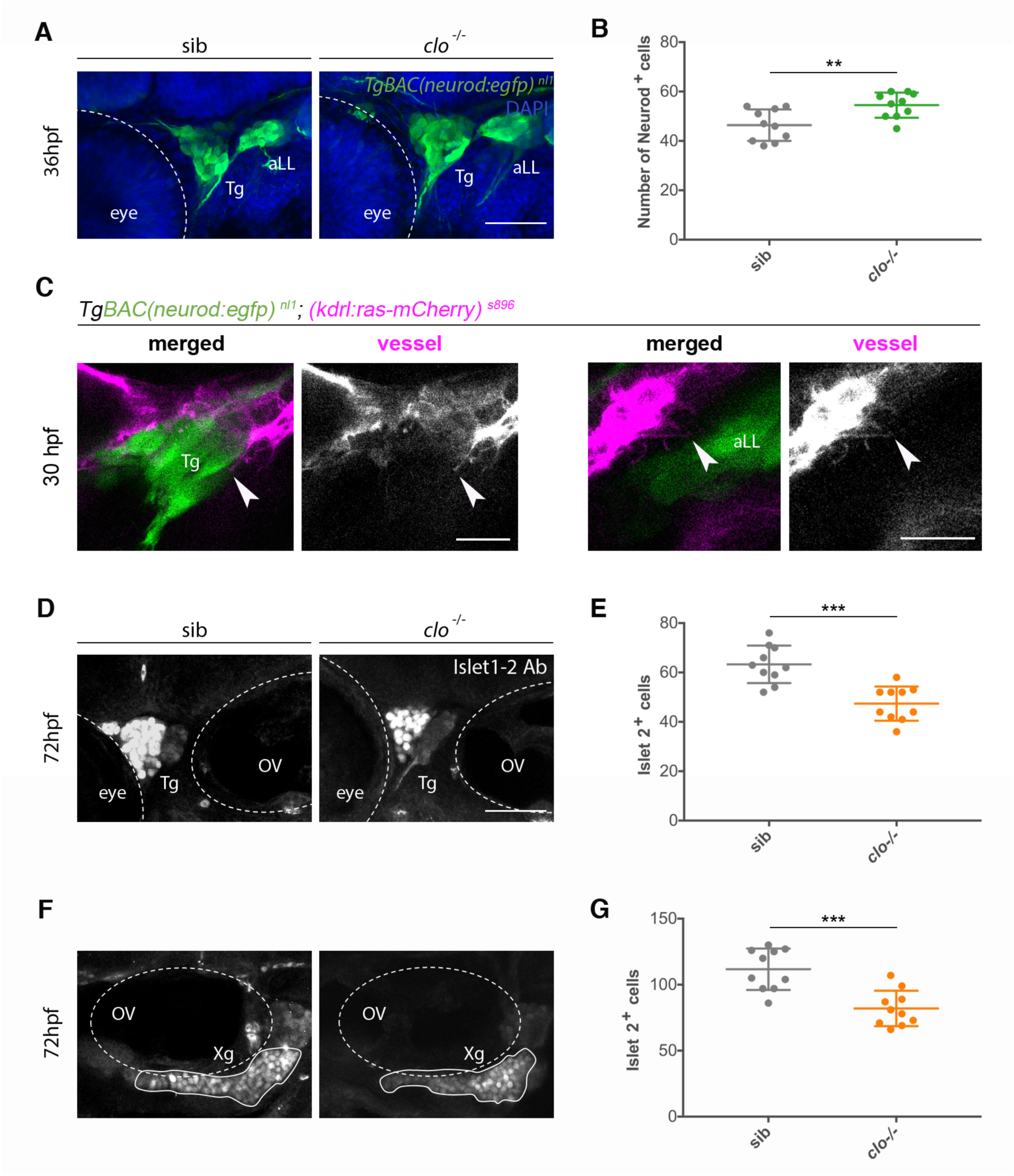
Filopodial contacts and defects on sensory neurogenesis in absence of vasculature in other sensory ganglia. (A) Confocal lateral images of Tg and aLL ganglia in sib and clo^-/-^ *TgBAC(neurod:egfp)* ^*nl1*^ embryos, at 36 hpf. Neurod^+^ cells: green, neuroblasts; DAPI: blue, nuclei. (B) Quantification of Neurod^+^ cells in the Tg of sib and clo^-/-^ embryos, at 36 hpf (n = 10). (C) Confocal lateral images of a single plane showing filopodial processes from the PHBC contacting neuroblasts of the Tg and aLL ganglia. Arrowheads indicate filopodia. (D-G) Confocal lateral images showing the number of Islet2^+^ nuclei in the Tg (c) and Xg ganglia (e) through Islet 1/2 antibody staining and quantified in E and G respectively (n = 10), at 72 hpf. Scale bars 50 µm for A, B, D-G, and 20 µm for C. Error bars, mean ± SEM. Unpaired two-tailed Student’s t-test. * p < 0.05, ** p < 0.01, *** p < 0.001. ns, non-significant. Tg, Trigeminal ganglion; aLL, anterior Lateral Line, OV, Otic Vesicle; Xg, Vagus ganglion; sib, siblings.

**Figure S5.**
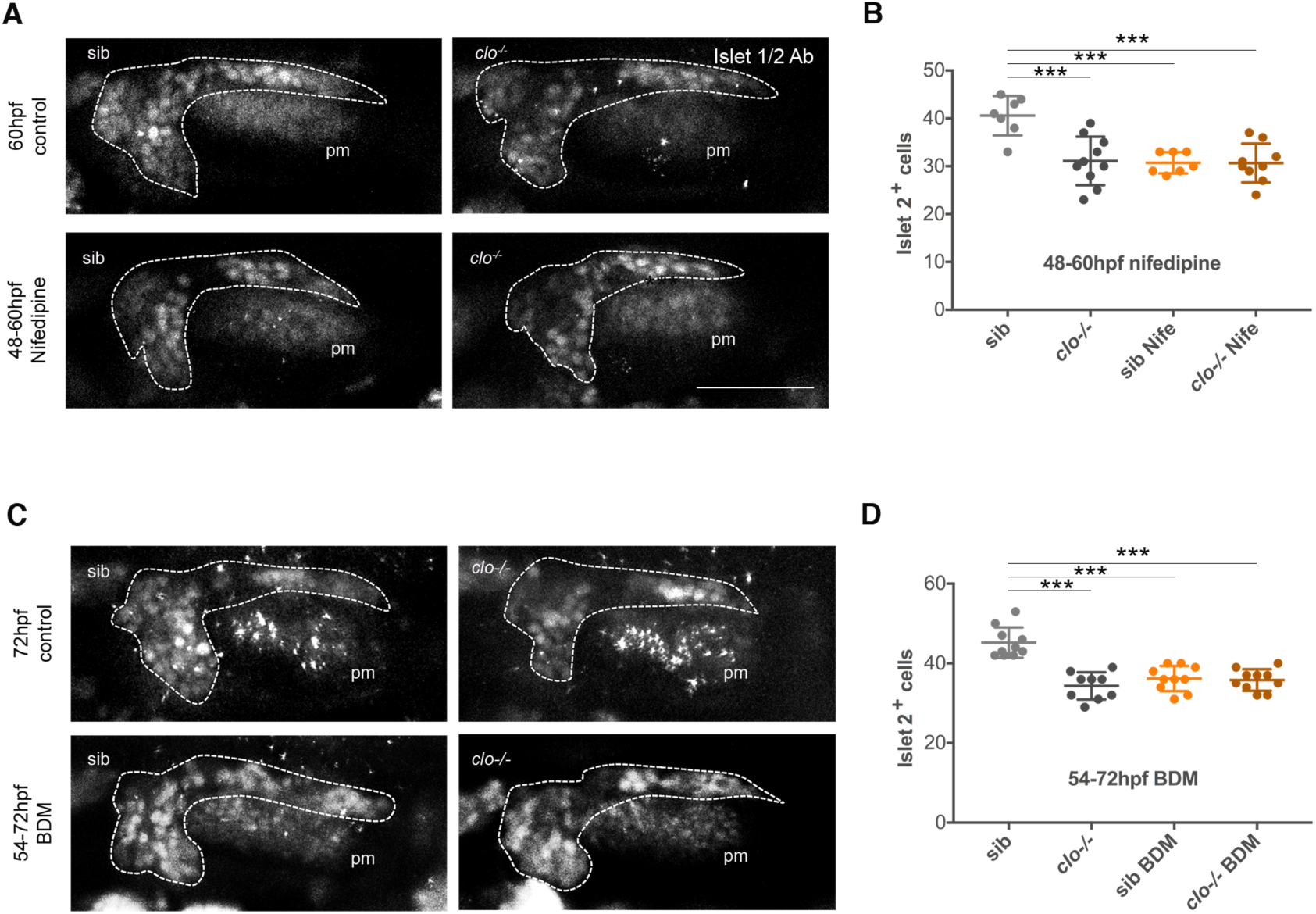
Blood flow arrest by Nifepidine at 60-72 hpf does not affect neuronal differentiation, while BDM at 54-72 hpf does. (A,B) Confocal images showing immunostained Islet2^**+**^ nuclei in the SAG (dotted line, visualized by *TgBAC(neurod:egfp)* ^*nl1*^) at 60 hpf in control (untreated) and 48-60 hpf Nifedipine treated sib and clo^**-/-**^ embryos; and quantified in I. 60 hpf sib (n = 7), 60 hpf clo^**-/-**^ (n = 10), 48-60 hpf sib (n = 7), 48-60 hpf clo^**-/-**^ (n = 9). (C) Confocal lateral images showing immunostained Islet2^+^ nuclei in the SAG (dotted line, visualized by *TgBAC(neurod:egfp)* ^*nl1*^) in sib and clo^-/-^ embryos at 72 hpf in control (untreated) and 54-72 hpf BDM treated embryos. (D) Graph showing the number of Islet 1/2^+^ cells in sib (n = 10), clo-/-(n = 9), 54-72 hpf sib BDM (n = 10), and 54-72 hpf clo^-/-^ BDM (n = 10). Scale bar 50 µm. Error bars mean ± SEM. Unpaired two-tailed Student’s t-test. * p < 0.05, ** p < 0.01, *** p < 0.001. ns, non-significant. pm, posterior macula; sib, siblings.

**Movie S1 Filopodial processes of the endothelial cells and neuroblasts contact each other at 30 hpf.**

Z-projection time-lapse video of *TgBAC(neurod:egfp)* ^*nl1*^; *(Kdrl:ras-mCherry)* ^s896^ embryos showing filopodia extending from the PHBC to the SAG and from the SAG to the PHBC at 30 hpf. Note thickenings in some neuroblasts filopodial. Lateral view. Scale bar 10 µm.

**Movie S2 Filopodial processes of the endothelial cells and neuroblasts contact each other at 30 hpf.**

Z-projection time-lapse video of *TgBAC(neurod:egfp)* ^*nl1*^; *(Kdrl:ras-mCherry)* ^s896^ embryos showing filopodia extending from the PHBC to the SAG and from the SAG to the PHBC at 30 hpf. Note dynamism of filopodial processes. Lateral view. Scale bar 10 µm.

**Movie S3 Filopodial processes of the endothelial cells and neuroblasts contact each other at 36 hpf.**

Z-projection time-lapse video of *TgBAC(neurod:egfp)* ^*nl1*^; *(Kdrl:ras-mCherry)* ^*s896*^ embryos showing filopodia extending from the PHBC to the SAG and from the SAG to the PHBC at 36 hpf. Note dynamics of filopodial processes. Lateral view. Scale bar 10 µm.

**Movie S4 Filopodial processes of the endothelial cells and neuroblasts contact each other at 36 hpf.**

Z-projection time-lapse video of *TgBAC(neurod:egfp)* ^*nl1*^; *(Kdrl:ras-mCherry)* ^*s896*^ embryos showing filopodia extending from the PHBC to the SAG and from the SAG to the PHBC at 36 hpf. Note length of EC filopodia. Lateral view. Scale bar 10 µm.

